# The Information Theory of Developmental Pruning: Optimizing Global Network Architecture Using Local Synaptic Rules

**DOI:** 10.1101/2020.11.30.403360

**Authors:** Carolin Scholl, Michael E. Rule, Matthias H. Hennig

## Abstract

During development, biological neural networks produce more synapses and neurons than needed. Many of these synapses and neurons are later removed in a process known as neural pruning. Why networks should initially be over-populated, and processes that determine which synapses and neurons are ultimately pruned, remains unclear. We study the mechanisms and significance of neural pruning in model neural network. In a deep Boltzmann machine model of sensory encoding, we find that (1) synaptic pruning is necessary to learn efficient network architectures that retain computationally-relevant connections, (2) pruning by synaptic weight alone does not optimize network size and (3) pruning based on a locally-available proxy for “sloppiness” based on Fisher Information allows the network to identify structurally important vs. unimportant connections and neurons. This locally-available measure of importance has a biological interpretation in terms of the correlations between presynaptic and postsynaptic neurons, and implies an efficient activity-driven pruning rule. Overall, we show how local activity-dependent synaptic pruning can solve the global problem of optimizing a network architecture. We relate these findings to biology as follows: (I) Synaptic over-production is necessary for activity-dependent connectivity optimization. (II) In networks that have more neurons than needed, cells compete for activity, and only the most important and selective neurons are retained. (III) Cells may also be pruned due to a loss of synapses on their axons. This occurs when the information they convey is not relevant to the target population.

## 1 Introduction

The number of neurons and synapses initially formed during brain development far exceeds those in the mature brain (Innocenti and Price, 2005). Up to half of the cells and connections are lost due to pruning (Huttenlocher et al., 1979; Huttenlocher and Dabholkar, 1997; Oppenheim, 1991; Stiles and Jernigan, 2010; Petanjek et al., 2011; Yuan and Yankner, 2000). This maturation process through initial over-growth and subsequent reduction suggests that the optimal wiring of the brain cannot be entirely predetermined. Instead, experience-dependent plasticity allows for a topological refinement of neural circuits, thereby adapting to the animal’s environment (Johnston, 2004). The removal of unneeded neurons and synapses reduces the high costs of the brain in terms of material and metabolism (Bullmore and Sporns, 2012). Pruning may also help to amplify neuronal signals against synaptic noise and support competition among synapses, thereby improving input selectivity (Neniskyte and Gross, 2017). Pathological over-pruning, however, may result in neuronal dysfunction, manifesting as cognitive impairments and clinical disorders such as schizophrenia (Feinberg, 1982; Johnston, 2004; Sekar et al., 2016; Sellgren et al., 2019). There seems to be a sweet spot between pruning and over-pruning for the brain to remain adaptive and resilient against damage.

What characterizes the cells and synapses that survive as opposed to the ones that die? A key factor for survival is neuronal activity. Initially, spontaneous activity is thought to drive survival. The refinement of cortical circuits then relies increasingly on sensory-driven and thus experience-dependent neuronal activity (Katz and Shatz, 1996). Neurons are thought to compete to activate target neurons in order to receive trophic supply (Oppenheim, 1989; Bonhoeffer, 1996; Lo, 1995). Although highly active presynaptic neurons are favored in this competition, the strengthening of connections also depends on coincident activity of the postsynaptic neuron. For instance, when postsynaptic cells in the primary visual cortex were pharmacologically inhibited, their more active afferents were weakened (Hata et al., 1999). Activity-dependent plasticity is thus based on bidirectional signaling between the presynaptic neuron and the postsynaptic neuron.

In this work, we employ Restricted Boltzmann Machines (RBMs) (Smolensky, 1986) as an artificial neural network model to study activity-dependent pruning. Similar to neuronal sensory systems, RBMs extract and encode the latent statistical causes of their inputs. Bernoulli RBMs consist of two layers of stochastic binary units, resembling the spiking of neurons. The visible units correspond to sensory input, while the hidden units encode a latent representation of this data. By adjusting their hidden representations as to maximize the likelihood of the data (Hinton, 2002), RBMs learn in an unsupervised fashion that resembles the Hebbian learning seen in neural networks (Whittington and Bogacz, 2017; Kermiche, 2019). RBMs have been used as statistical models of sensory coding (Zanotto et al., 2017; Gardella et al., 2018; Rule et al., 2020). Furthermore, multiple RBMs can be stacked to build a Deep Boltzmann Machine (DBM) (Hinton and Salakhutdinov, 2006) to model the hierarchical, bidirectional computation of the visual system (Hochstein and Ahissar, 2002; Turcsany et al., 2014). Such DBM models have been used, e.g., to explore how loss of input could lead to visual hallucinations in Charles Bonnet Syndrome (Reichert et al., 2013).

Here, we explore several local and biologically plausible rules for iteratively pruning unimportant synapses and units from DBM networks. Many models of neural pruning simply remove small synaptic weights (Chechik et al., 1998; Han et al., 2015; Mimura et al., 2003). However, magnitude need not equal importance, and small weights can be necessary to maintain a small error (Hassibi et al., 1993). A more theoretically grounded measure of synapse importance may lie in the activity of individual neurons and their correlations (Iglesias and Villa, 2007).

Information-theoretic approaches to network reduction provide a principled starting point. For example, the Optimal Brain Surgeon algorithm (LeCun et al., 1990) estimates each parameter’s importance by perturbing its value and re-evaluating an error function: low changes in error indicate redundant, uninformative parameters. This curvature with respect to small parameter changes is given by the Hessian of the error function, and is locally approximated by its negative, the Fisher Information Matrix (FIM). To reduce computational complexity, Optimal Brain Damage (as opposed to Optimal Brain Surgeon (Hassibi et al., 1993; Hassibi and Stork, 1993; Dong et al., 2017)) makes the simplifying assumption of a diagonal Hessian matrix.

Fisher Information (FI) based estimates of parameter importance have recently been used to overcome catastrophic forgetting in artificial neural networks (Kirkpatrick et al., 2017). In contrast to identifying parameters worth keeping, we aim to identify parameters worth removing. For RBMs, the diagonal of the FIM can be estimated locally based on the firing rates of units and their coincidence (Rule et al., 2020; Deistler et al., 2018). In principle, this estimate of parameter importance is available to individual synapses and neurons, and could thus guide the search for efficient representations during neurodevelopmental pruning.

We organize the results as follows: first, we introduce our estimates of synaptic importance based on locally-available activity statistics. Next, we discuss the overall network curvature and demonstrate that important synapses centre on overall highly informative neurons. Based on these observations, we introduce local pruning rules to iteratively remove synapses and neurons from RBMs and DBMs that were trained on image patches from two different data sets. We evaluate the fit of the pruned models across different pruning criteria by assessing their generative and encoding performance. Finally, we discuss the biological plausibility of our activity-dependent pruning rules by comparing them to alternative rules used in our own and related work and provide implications of our findings.

## 2 Results

### 2.1 An activity-dependent estimate of Fisher Information

The goal of this work is to derive and use a local pruning rule to reduce network size and identify a relevant network topology in RBMs and DBMs. By relevant network topology we mean a topology that closely matches computation, in the sense that it only includes neurons and synapses that are relevant for the task at hand. In our experiments this task is the encoding of visual stimuli in hidden representations of RBMs and DBMs. RBMs are energy-based models whose energy function is given by:

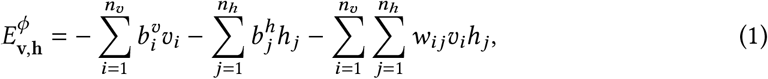

where *v*_*i*_ stands for visible unit *i*, *h*_*j*_ stands for hidden unit *j* and *w*_*ij*_ for the bidirectional weight connecting them. The biases 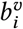 and 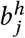 correspond to the excitability of a neuron. All weights and biases make up the parameter set *ϕ* = {**W, b^v^, h^h^**}. As we work with Bernoulli RBMs, all neurons are binary: they are either firing (1) or silent (0).

The Hessian of the objective function of a model gives information about the importance of parameters and can be used for pruning (LeCun et al., 1990; Hassibi et al., 1993). It can be locally approximated by its negative, the FIM. However, for FI to be a suitable indicator for self-relevance to biological neurons, it must also be available locally at each synapse. This rules out using the full FIM, since this requires information about the connectivity of the whole network. Presumably, neurons in the brain use a local heuristic that can be computed based on information available to single synapses. It was recently shown that local activity statistics indeed correlate with importance as assessed by FI (Rule et al., 2020). In the case of the RBM this correspondence is exact; an entry of the FIM for an RBM has the form (Rule et al., 2020):

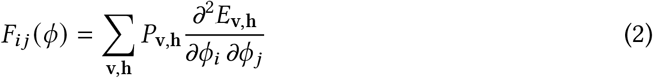

The FIM locally approximates the Kullback–Leibler divergence between the model distribution under the current parameter set *ϕ* and the model distribution when a pair of parameters *ϕ*_*i*_ and *ϕ*_*j*_ is slightly perturbed. When *F*_*ij*_ tends towards zero, the two parameters *ϕ*_*i*_ and *ϕ*_*j*_ are redundant. Through sampling we can average over *P*_*v,h*_. For instance, for two weights *w*_*ij*_ and *w*_*kj*_ connecting a presynaptic to a postsynaptic neuron, respectively, we get (Rule et al., 2020):

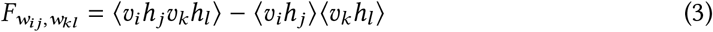

The resulting entries of the FIM depend on coincident firing of pre- and postsynaptic neurons and are arguably locally available.

### 2.2 Large encoding models have many poorly specified parameters

In a first line of experiments, we inspected the curvature of the energy landscape of overcomplete models that had more latent units than needed to encode the visible stimuli. We started by fitting overcomplete RBMs to circular, binarized patches sliced out of images of natural scenes (see Figure 1 A). Parameter-wise FI was computed by our activity-dependent estimate of FI from Equation 2 (see Appendix A.1 and (Rule et al., 2020)). For these relatively small models, we computed the full FIM, which typically turned out to be sparse (see Figure 1 B). This indicates that large encoding models indeed have many poorly specified parameters, which can be safely removed from the model.

**Figure 1:**
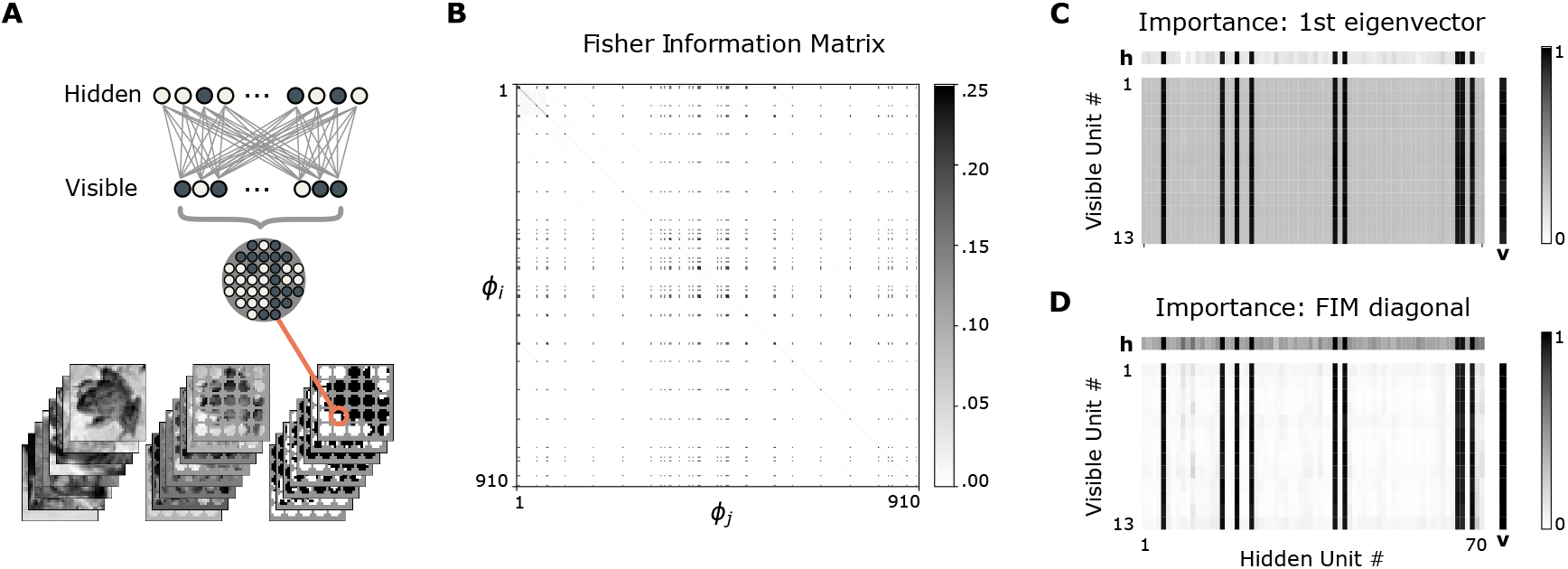
**(A):** Preparation of dataset and general model architecture: circles of different radii were sliced out of CIFAR-10 images and binarized. The pixels corresponded to the visible units of a standard RBM with one hidden layer. **(B):** FI for an exemplary over-parameterized RBM initialized with *n*_*v*_ = 13 visible units and *n*_*h*_ = 70 hidden units. The FIM is sparse, indicating many irrelevant parameters. **(C):** Normalized parameter importance estimated from the first eigenvector of the FIM, separately for the weights (matrix), the hidden biases (horizontal), and visible biases (vertical) **(D):** Normalized parameter importance directly estimated from the diagonal entries of the FIM.

The sparseness of parameter-wise FI is also evident from a visualization of the first eigen-vector and the diagonal of the FIM, separately for the weights, hidden biases and visible biases (see Figure 1 C and D). Furthermore, the eigendecomposition of the FIM revealed that the largest eigenvalue is larger than the second largest eigenvalue by an order of magnitude (first eigenvalue 23.97, second eigenvalue 1.60).

The correspondence between parameter importance estimated from the first eigenvector and from the diagonal supports our use of Optimal Brain Damage for larger models, when computing the full FIM was no longer feasible. Strikingly, the important weights typically aligned with few hidden units and their biases. This structure of the FIM suggests a separation into important hidden units and unimportant ones. It follows that FI motivated pruning likely leads to entire units becoming disconnected, which would allow their removal from the network. This would correspond to neuron apoptosis after excessive synaptic pruning.

Overall, these empirical investigations reveal that there are only few overall important units, and that important weights align with them. This confirms that parameter importance as revealed by the FIM contains meaningful information for pruning, and that synapse pruning can eventually lead to the pruning of whole neurons that lack relevant afferent or efferent connections.

### 2.3 Local pruning rules based on parameter-wise Fisher Information

We now present our results of applying the local estimate of FI we introduced above as a criterion to prune RBMs. We started by removing poorly specified, unimportant weights according to different pruning criteria. For a pruning criterion based on the full FIM, we used the weight-specific entries of its first eigenvector as a direct indicator of weight importance. Next, we used our local estimate, where we considered the FIM diagonal only, as suggested by Optimal Brain Damage.

Computing the diagonal entries of the FIM corresponds to Equation 3, when the two modified parameter values are the same (*w*_*ij*_ = *w*_*kl*_). It captures the effect on the error when we modify a single parameter instead of a parameter pair at a time. For the FI of the weights, the equation thus simplifies to (Rule et al., 2020):

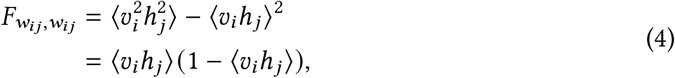

where 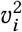 and 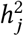 simplify to *v*_*i*_ and *h*_*j*_, respectively, because neural activation is binary.

In this scheme, weight importance is estimated from the covariance of a pre- and post-synaptic neuronal firing. This variance estimate of FI is both activity-dependent and locally available, and is therefore a biologically plausible indicator of synapse importance. However, neurons need not track this covariance explicitly. Activity-driven pre-post correlations influence synaptic weights and vice versa. As a result, statistical quantities relevant to pruning can be encoded in the synaptic weights themselves. In Appendix A.2 we derive a mean-field approximation of the pre-post correlations that depends only on the synaptic weight and the mean firing rates of the presynaptic and postsynaptic neuron. This offers a proof of principle that locally-available quantities can provide the signals needed for a synapse (and by extension a neuron) to assess its own importance in the network. We refer to this alternative FI-based pruning rule as the heuristic FI estimate (as opposed to the variance FI estimate).

For our synaptic pruning experiments, we ranked weights according to their importance assessed by one of the aforementioned local estimates, or by the first eigenvector of the FIM. We then iteratively deleted half of the weights with lowest estimated importance, while monitoring model fit. Unconnected hidden units were removed from the model. Absolute weight magnitude, random pruning and removal of the *most* important weights (“Anti-FI” pruning) served as reference criteria for pruning. Figure 2 shows the model’s generative performance over the course of three pruning iterations, evaluated immediately after pruning and after allowing for retraining.

**Figure 2:**
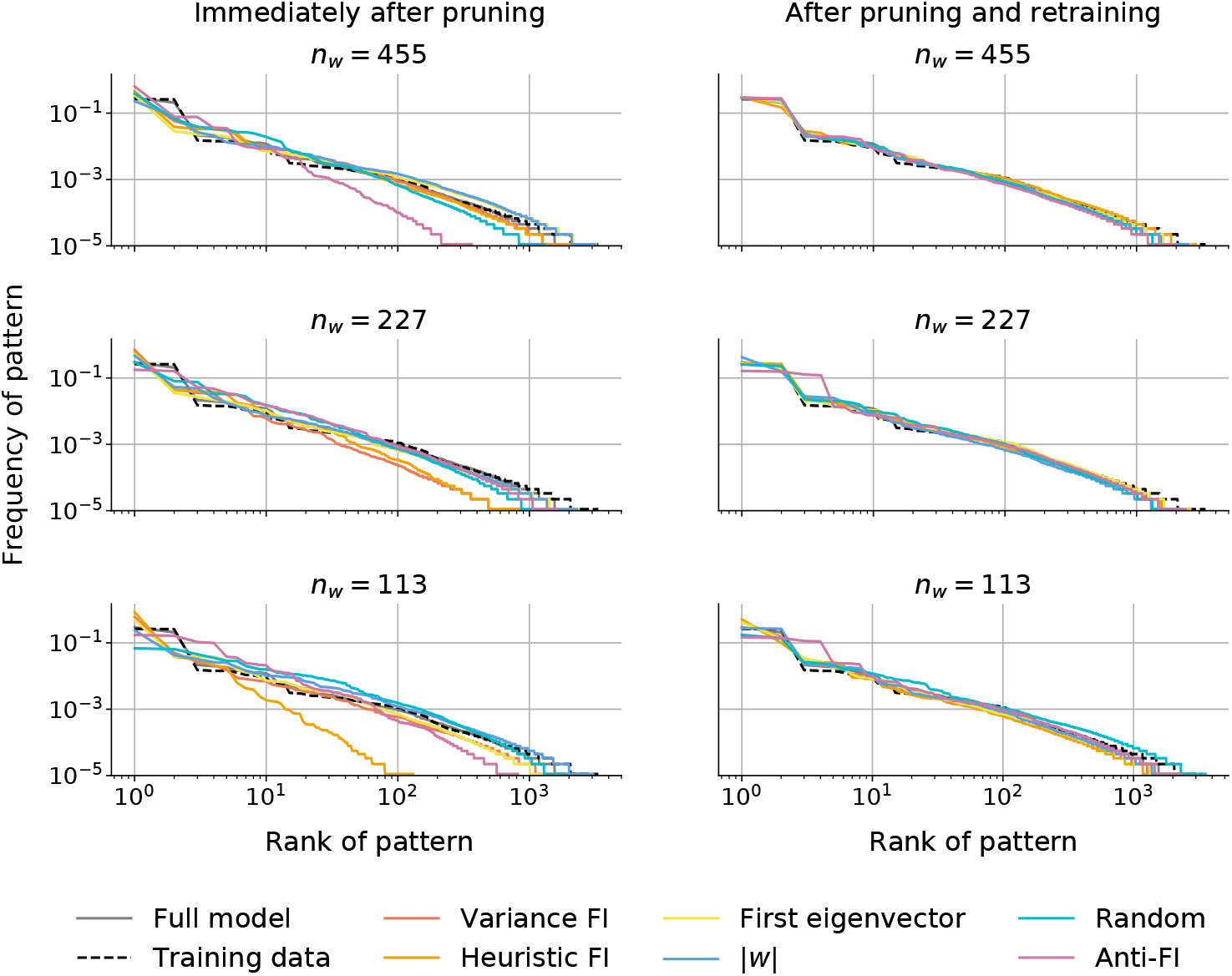
Rank-frequency plots for generated patterns by a pruned RBM which initially had *n*_*v*_ = 13, *n*_*h*_ = 70, and *n*_*w*_ = *n*_*v*_ × *n*_*h*_ = 910 weights. Half of the weights were removed in each of three pruning iterations. For all pruning strategies except Anti-FI pruning the model retains the ability to match the distribution of the training data (dashed lines) after retraining, indicating good generative performance.

Generally, excessive pruning was detrimental to generative performance, yet retraining repeatedly rescued the pruned model: with moderate amounts of pruning, training restored network performance irrespective of the pruning criterion. In biological neural networks, on-going training occurs continuously alongside activity-driven pruning, so a period of retraining following pruning is plausible. The network performance immediately following pruning differed depending on the pruning criterion and the number of remaining weights. Performance remained relatively stable in the first iteration of pruning, but was adversely affected later: the smaller the model, the more it suffered from pruning. With decreasing model size the effect of different pruning criteria became more pronounced, as we shall show next. Anti-FI pruning clearly damaged the network’s generative performance even in the first iteration of pruning, especially for rare patterns. Interestingly, random pruning barely affected the generative performance.

A reason for this may lie in the number of hidden units that became disconnected through pruning. Table 1 compares the number of remaining hidden units *n*_*h*_ against the number of remaining weights *n*_*w*_, separately for each pruning criterion. While the number of remaining weights *n*_*w*_ after each pruning event was kept equal across pruning criteria, there was a great difference in the number of units that became disconnected from repeated weight pruning. FI typically concentrated on few hidden units (see Figure 1), contrasting them with units of low overall importance. Such uninformative units were left disconnected after repeated pruning iterations. The sudden loss of many units damaged the model’s generative performance immediately after pruning, but performance recovered with retraining. In contrast, random pruning uniformly deleted weights from all units such that only few became disconnected. Pruning the largest weights (weight magnitude) and most important weights (Anti-FI) also kept more hidden units in the model, making the final encoding less efficient than the one achieved through FI-motivated pruning.

**Table 1:**
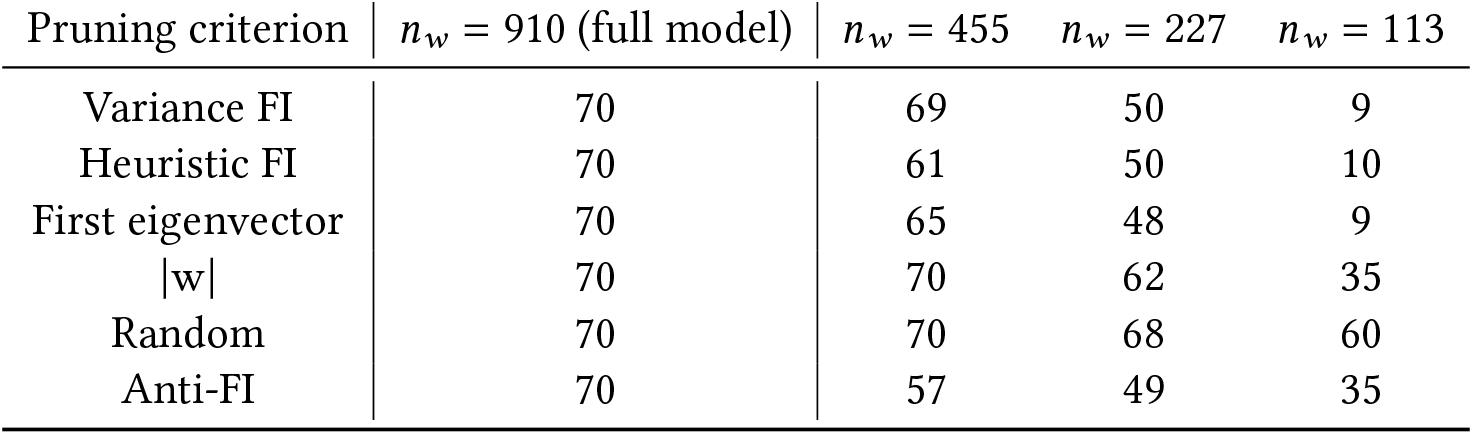
Differences in numbers of remaining hidden units *n*_*h*_ in the RBM. During each of the three pruning iterations half of the weights were removed according to different criteria. *n*_*w*_ describes the remaining weights at a time. The cells contain the number of neurons after each pruning iteration.

In sum, for all pruning strategies (except Anti-FI pruning, which removed a large fraction of important weights), the network could recover from the loss of weights and units through retraining (Liu et al., 2018; Crowley et al., 2018). However, the resulting latent representations differed: the number of disconnected neurons varies strongly between the pruning methods. This in turn may affect subsequent read-out from these representations, an effect we will investigate next using deep networks.

### 2.4 FI-guided pruning retains useful deep encoding

Above, we demonstrated an iterative approach of pruning weights from RBMs while monitoring their fit from the frequency of its generated patterns. In line with results by (Liu et al., 2018; Crowley et al., 2018), pruned networks generally required retraining. In small RBMs that encoded simple visual patterns, retraining could always compensate for the loss of synapses, regardless of the pruning criterion. However, we found that FI pruning, as opposed to pruning small weights, can reduce the cost of the network in terms of the number of remaining neurons. This supports the view that the function of pruning is to find an appropriate architecture, rather than specific parameter configurations.

To investigate the effect of the pruning method on the latent representation, we increased the complexity of the model architecture by adding another hidden layer, resulting in a DBM. We trained this multi-layer model on a labeled dataset to better quantify the fit of the model and quality of the latent representation during iterative pruning.

Specifically, we used the MNIST handwritten digits dataset (LeCun et al., 2010), and tested both the generative and classification performance using the latent activations. To measure encoding quality, we compared the accuracy of a logistic classifier trained on the stimulus-evoked activity in the latent units (“encodings”) to one trained on the raw digits. To evaluate the generative performance, we asked a classifier trained on the raw digits to categorize samples and predict probabilities of them belonging to each digit class. These were summarised as digit-wise quality scores and diversity scores for the generated patterns (see Methods).

Each image of the dataset was binarized and scaled to 20×20 pixels. To simplify computation and add biological realism, we restricted the receptive fields of each unit in first hidden layer **h**^1^ to a small region of the visible inputs. To encode digits, the network was therefore forced to combine these lower-level features in the second hidden layer **h**^2^ (see Figure 3 A). We then assessed encoding quality based on the accuracy of a classifier applied to the stimulus-driven activity in the deepest hidden network layer. Figure 3 B shows the classification errors over the course of 10 pruning iterations according to different criteria. Before pruning, the classifier performed better using the latent states of the trained network compared to the raw digits.

**Figure 3:**
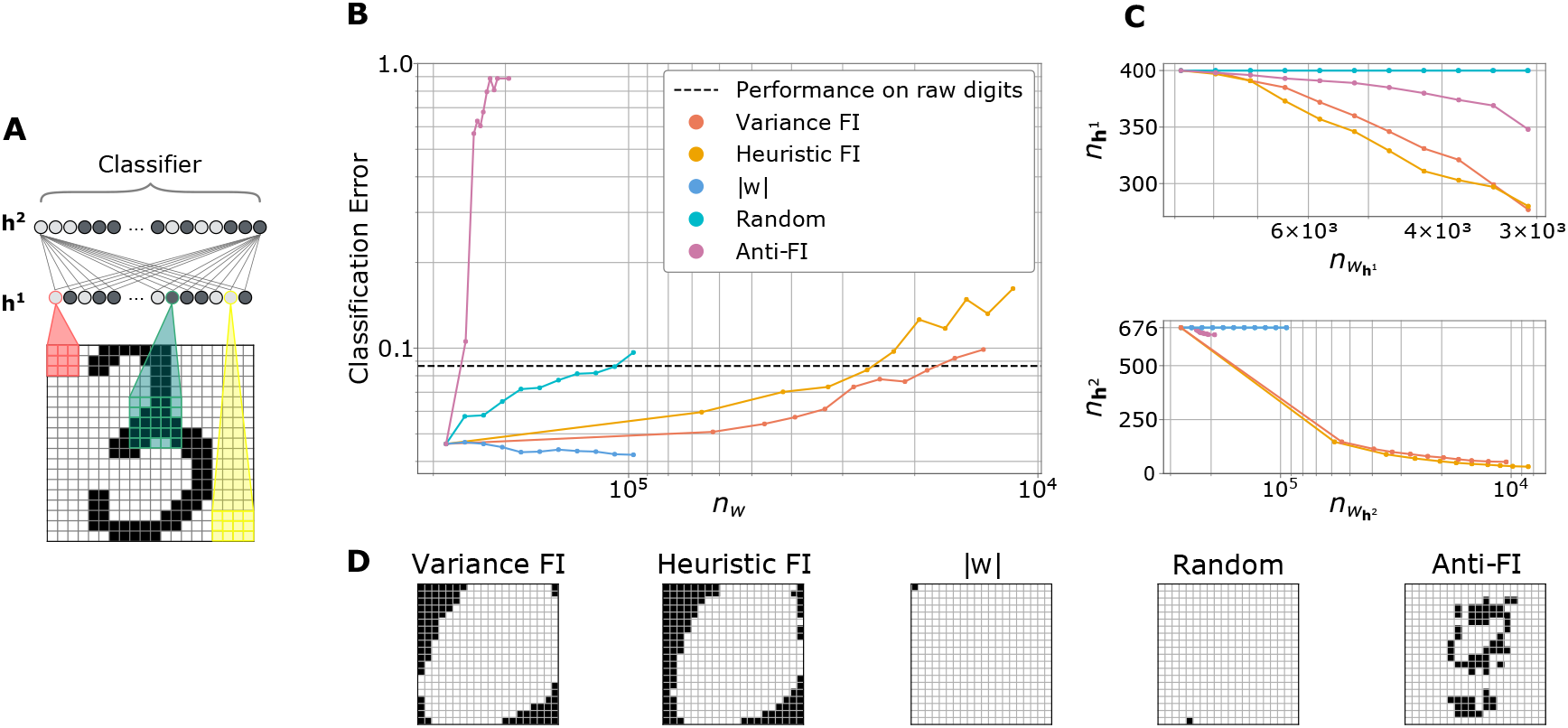
**(A):** Data representation and DBM architecture. MNIST digits were cropped and binarized. Each unit from hidden layer **h**^1^ had an individual receptive field covering neighboring input pixels; **h**^2^ was fully connected. Only few connections are shown due to clarity. The classifier was trained on latent encodings from **h**^2^. **(B):** Classification error of a logistic regression classifier trained on **h**^2^ samples as a function of remaining weights *n*_*w*_. The dotted line stands for the baseline error of a classifier trained on the raw digits. **(C):** Number of latent units in **h**^1^ and **h**^2^ as a function of remaining weights over the course of ten iterations of pruning. **(D):** Final visible layers after pruning according to different criteria. Black pixels denote disconnected units.

On each iteration, the 10th percentile of weight-specific FI or absolute weight magnitude was computed. In practice, more than 10% of FI estimates were often zero. In this scenario, we instead pruned all synapses with zero importance. Units of the intermediate layer **h**^1^ that became disconnected from either side (v or **h**^2^), were completely removed. Thus, the total number of deleted weights may be larger than specified by the importance measure as a secondary effect.

Similarly to the results for the single-layer RBM, pruning a modest number of DBM parameters and retraining did not have a dramatic effect on the latent encoding, as measured by the classification error (Figure 3 B). Performance usually decreased immediately after pruning (not shown), but could be compensated for through short retraining. With successive pruning iterations, performance decreased steadily. Interestingly, pruning according to absolute weight magnitude retained a useful encoding for the classifier; its error even decreased in the course of ten pruning iterations. Yet the effective number of weights removed in this case was small, and very few units were disconnected.

In contrast, removing randomly selected weights led to rapid degradation, and removing important weights (Anti-FI pruning) was particularly harmful: classification performance deteriorated after the first pruning event, and remained below chance even with retraining. This suggests that Anti-FI pruning causes a loss topologically relevant neurons. Visualizing the pixels corresponding to the remaining units in the visible layer v after 10 pruning iterations in Figure 3 D shows this clearly: While FI-based pruning correctly disconnects pixels at the boundaries where images contain no information, Anti-FI pruning removes links to the central pixels most useful for digit classification.

Figure 3 C shows the number of remaining units as a function of remaining weights in the two latent layers of the DBMs. The difference in the number of lost units agrees with with what we observed for RBMs (see Table 1). FI-guided pruning allowed for the removal of units in all layers of the model, thereby making the encoding more efficient. The encoding also remained useful for classification. When weights were pruned according to both our estimates of FI, the performance of the classifier only fell below baseline after the number of weights was reduced by more than one order of magnitude. Yet neither random pruning nor pruning by absolute weight magnitude led to any of the hidden units becoming disconnected. In the visible layer, only one pixel became disconnected in each case (see Figure 3 D). Pruning according to our estimates of parameter-wise FI on the other hand left a number of hidden units as well as uninformative visible units at the borders of the images unconnected.

Taken together, these results show that the pruning criterion matters in deeper networks. The network performance can either recover through retraining if activity-dependent pruning is used (FI pruning and pruning by weight magnitude) or it is permanently damaged in case of random pruning and Anti-FI pruning. Furthermore, FI-pruning produces the most efficient network topology as this allows selectively the retention of the most important neurons, unlike simpler strategies such as pruning by weight magnitude.

### 2.5 Models that lost informative synapses and neurons can no longer generate meaningful patterns

In addition to classification accuracy, it is relevant to assess how pruning affects the generative performance of the networks, since this measures how well they capture the features of the input data. Previous studies have explored DBMs as a generative model for some types of hallucinations, for example those seen in Charles Bonnet Syndrome (Reichert et al., 2013). In this disease, patients experience vivid hallucinations after going blind. DBMs may be a suitable model for this because they are probabilistic generative models: without supervision, they learn a statistical representation in the hidden layers that can be used to generate the visible samples. To simulate the loss of vision, Reichert et al. (2013) completely removed the visible layer of a DBM. Due to homeostatic compensation, spontaneous activity in the hidden units continued to reflect plausible visual inputs, which might be interpreted as hallucinations by downstream readouts. Inspired by these results, we investigated the generative performance of partly lesioned DBMs over the course of pruning.

To quantify generative performance, we provided samples of visible patterns from the pruned models to a classifier trained on the raw images of the digits. We used the classification confidence to measure digit quality (Figure 4 A). When the model was deprived of its most informative weights (Anti-FI pruning), retraining could not compensate for the loss of relevant weights. A much weaker, but significant degradation was also observed for random pruning. In contrast, the quality of generated digits by DBMs pruned by weight magnitude, or using the FI-based rules, suffered less after allowing for retraining. Even though far more weights were removed, the models still produced recognizable digits (Figure 4B).

**Figure 4:**
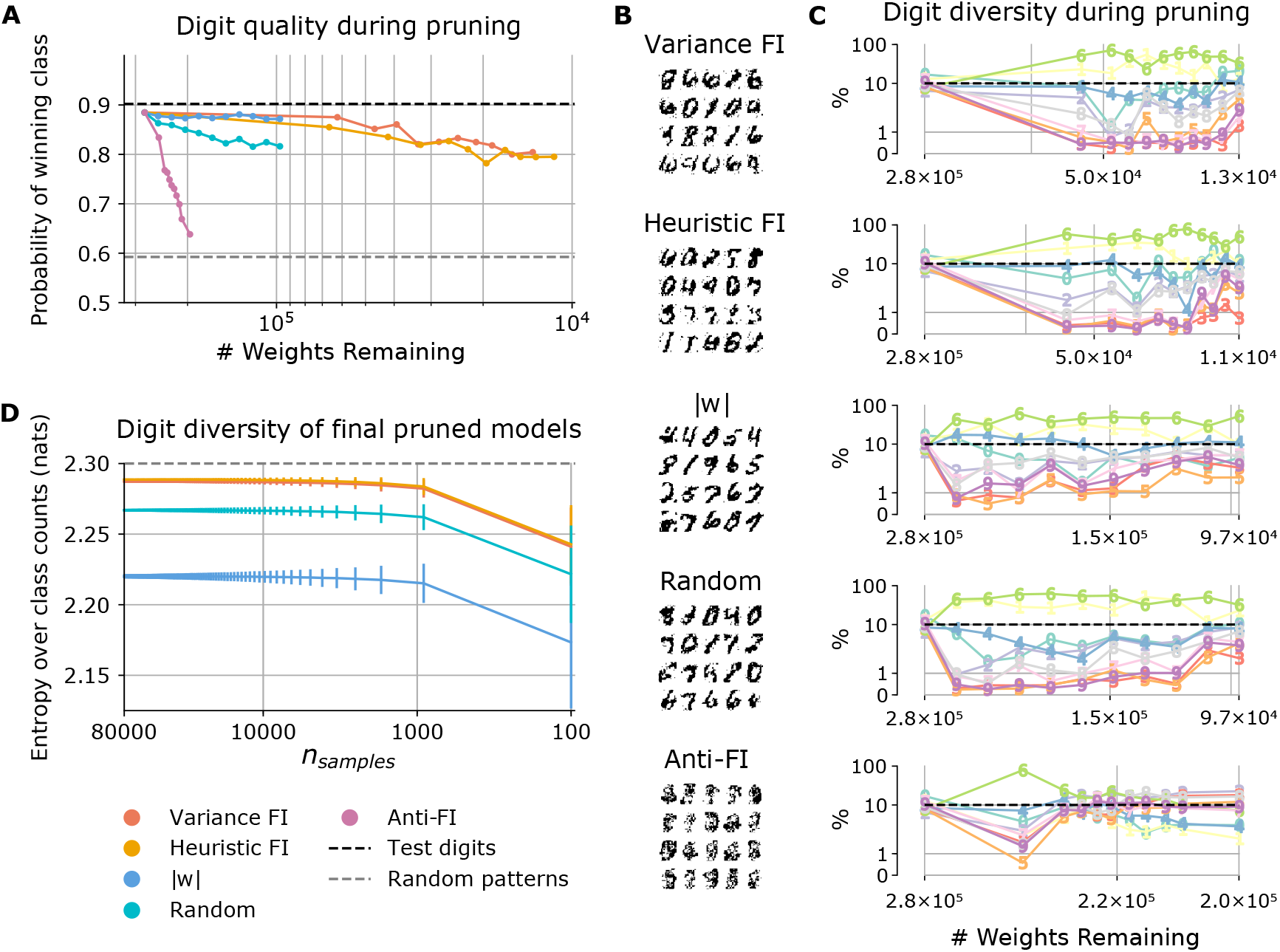
**(A):** Maximum class probability assigned to generated samples from pruned networks. This summarizes the confidence of a classifier that the generated digits belonged to a specific class. The back and gray dashed line show the confidence of the classifier for the MNIST test digits and randomly generated patterns, respectively. **(B):** Examples of generated patterns after ten iterations of pruning. **(C):** Percentage of samples that were assigned to one of 10 color-coded digit classes as a function of remaining weights. X-axis limits represent initial and final numbers of weights. Dotted line indicates completely balanced samples at 10%. Pruning altered the distribution of generated patterns, perhaps because some number shapes were more central than others in the classifier’s model of digit shapes. **(D):** Entropy over the distribution of generated digits, computed from the generated samples after retraining the fully pruned networks from scratch, with random weight initialization. The small variations of the entropies for larger sample sizes indicate unbiased estimates for the largest sample used here. An entropy value of ≈ 2.30 nats corresponds to even coverage of all digits. The plot shows the mean entropy computed by bootstrapping: 10,000 samples of size k were were drawn (with replacement) from the generated digits. Error bars indicate one standard deviation.

Finally, we compared the diversity of the generated digits (Figure 4 C). We used a classifier to categorize each sample into one of the ten digits. In the un-pruned model, the fraction of samples categorized as one of each of the ten digits ranged from from 6% to 17%, indicating good generative performance (if the samples were completely balanced, each digit would appear with a probability of 10%). Pruning with limited retraining increased class imbalance, e.g. the digit six was over-represented in all cases. Note, however, that this assessment of generative performance is not meaningful when the network generated degraded digits that were hard to classify, as was the case for Anti-FI pruning (compare examples in Figure 4 B).

These results suggest that pruning impairs the generative capabilities of the networks, either degrading the generated representations or causing a bias towards certain patterns. To test if this was a consequence of the pruned weight values verses network architecture, we fully retrained the pruned networks with random parameter initialization (Crowley et al., 2018). The evaluation of patterns generated by these networks shows that class balance can be largely restored, in particular in the FI pruned networks (Figure 4 D), indicating that the loss of digit diversity was not an architectural deficit.

To summarize these results, we computed the entropy of the generated digits distributions for different sample sizes, and linearly extrapolated the true entropy (Figure 4 D). The entropy of the FI-pruned models is close to the maximum value of 2.30 nats for fully balanced classes (variance FI: 2.287 nats, heuristic FI: 2.289 nats). Interestingly, while random pruning yields a lower entropy and thus higher class imbalance (2.267 nats), pruning by weight magnitude produces the most imbalanced network (2.22 nats). We excluded Anti-Fi pruning from this analysis, since the classifier confidence was too low for the class assignment to be meaningful.

Taken together, all FI pruning approaches preserved the generative quality to a similar degree. In contrast, the Anti-FI pruning algorithm disconnected relevant visible and hidden units, leading both sensory “blindness” and an inability of the model to meaningfully report the visual correlates of latent activity. Weight magnitude alone is an insufficient indication of parameter importance, and while networks pruned in this way still can be used to classify the digits well, they do no longer fully capture the statistics of the input images, and their “hallucinations” are less diverse than those of models pruned according to FI estimates. This suggests that to preserve the activity statistics in networks, pruning requires an activity-dependent mechanism.

## 3 Discussion

In this work, we used stochastic latent-variable models of sensory encoding to derive new theoretical results on neural pruning, and explored these results through numerical simulations. By examining the energy-based interpretation of such models, we showed that the importance of synapses and neurons can be computed from the statistics of local activity. To our knowledge, our work is the first empirical study of network reduction guided by an activity-dependent estimate of parameter-wise Fisher information in this context. Our pruning rule operates at single synapses, and provides a principled route to eliminating redundant units. Searching for biological analogues of these quantities and processes may provide insight into naturally occurring pruning in the developing brain.

### 3.1 FI-guided pruning identifies efficient network architectures

In biology, over-production of neurons and synapses likely confers advantages. These extra neurons could accelerate initial learning (Raman et al., 2019), or allow the network to identify better architectures than it would otherwise (Steinberg et al., 2020). After this initial learning, neurons and synapses not involved in covariant experience-dependent activity are lost (Katz and Shatz, 1996; Hata et al., 1999). Our theoretical derivations make this precise, deriving local pruning rules that estimate the importance of a connection from the covariant activity of a pre- and postsynaptic neuron. This captures a central intuition about importance: neurons that vary little are less relevant, and synapses with weak pre-post correlations are less important.

Pruning’s role in identifying efficient topologies is highlighted by the effect of our Anti-FI pruning rule, which removes the most important synapses first. On a network encoding MNIST handwritten digits, Anti-FI pruning removes units that are vital for performing the computation. In contrast, FI-based learning rules retain these important components. In this case, it is unsurprising that pruning removes irrelevant units in the visual periphery. However, parameter importance is less obvious in deeper layers of the network, and more generally for other stimuli and tasks.

It would also be worth studying the time-course of pruning and retraining, to see if artificial neural networks can reproduce the critical periods observed in biology. A recent study of the effects of perturbing the input during different time points of training in neural networks suggests that a critical learning period may be visible in a plateau of the FIM trace (Achille et al., 2018). It would be interesting to examine if such critical periods exist not just for changes in the “software” (inputs and weights) of a network, but also for its “hardware” (its topology and number of parameters).

### 3.2 Relationship to pruning in artificial neural networks

State-of-the-art models in machine learning are often over-parameterized, and optimizing such models via pruning is a subject of active research. The vast reduction of the number of neurons in the hidden layers in our FI-pruned models show how pruning could play a similar role in architecture optimization in the brain.

Yet some studies call into question the usefulness of pruning. It can be difficult to continue training networks with pruned weights without incurring performance losses (Crowley et al., 2018; Frankle and Carbin, 2018; Liu et al., 2018), as networks become trapped in local minima. It has thus been argued that the value of pruning is rather to find a suitable architecture for random re-initialization (Crowley et al., 2018; Liu et al., 2018) or well-initialized weights of sub-networks within larger networks (Frankle and Carbin, 2018). Both of these algorithms, however, cannot by implemented by biological neural networks because they include a reset of synaptic weights to previous values. In our work, incremental retraining of trained parameter values was sufficient to preserve performance. This is consistent with the strategy that must be employed in the brain, in which learning and pruning occur concurrently. It remains to be seen whether the brain is also subject to the trap of local minima, or whether additional mechanisms make the issue irrelevant in biological networks.

In addition to removing parameters, some machine learning algorithms also *add* units, to allow the network to adapt its size to the problem domain. Berglund et al. (2015) used a mutual-information based measure to both remove and add neurons in Boltzmann machines. It would be interesting to explore whether local information-theoretic rules could be extended to include activity-driven synaptogenesis and neurogenesis, as is observed in brain areas such as the the hippocampus (Kempermann, 2002). Potentially, our local estimate of FI could not only guide network reduction but also network growth, resulting in an integrative model of neuronal plasticity.

### 3.3 The role of ongoing learning and homeostasis

In biology, pruning and learning occur simultaneously. To emulate this, we retrained networks for ten epochs after each batch of pruning. This batched form of sensory-driven plasticity allowed the networks to recover from a performance drop observed immediately after pruning. Yet in biology, there are numerous types of homeostatic plasticity that could compensate for loss of cells and synapses without error-driven retraining (Wu et al., 2020). Synaptic depression and spike-frequency adaptation are examples for such pre- and postsynaptic adaptation mechanisms, respectively (Benda and Herz, 2003; Jones and Westbrook, 1996). Weight re-scaling resembling such adaptation mechanisms could potentially help make pruning less disruptive. Such homeostatic rules may accelerate learning by reducing the amount of training needed to compensate for the effects of pruning.

### 3.4 Pathological pruning, hallucinations, and neurological disorders

Model networks can provide insight into neurological disorders. For example, Reichert et al. (2013) use DBMs to model Charles Bonnet syndrome. In their model, homeostasis re-activated internal representations in “blinded” DBMs, creating hallucinations. Here, we explored how networks degrade when pruning is inaccurate or excessive. Unlike in Reichert et al. (2013), our networks continued to receive input. However, aberrant pruning led to partial blindness and neuronal loss. On the MNIST dataset, all pruned models (except the Anti-FI network) could still generate meaningful digit-like patterns, indicating that the basic circuits of perception were intact. However, pruning degraded the quality and diversity of generated digits. In random and weight-based pruning, digit generation was impaired even after complete retraining. FI pruning, in contrast, retained generative performance in comparatively smaller networks.

Statistical theories view hallucinations as errors in internal predictive models (Adams et al., 2016; Fletcher and Frith, 2009). Hallucinated stimuli in the absence of input is one form of this. Over-confidence in erroneous internal models, or incorrect predictions of which stimuli are likely, can also lead to mistaken —or hallucinatory— perception. Loss of generative diversity in our simulations indicates that the model’s internal priors no longer match the external world. Interpreted in the context of predictive coding, this bias implies that some stimuli are poorly predicted by internal states, and would therefore register as unexpected or surprising.

This suggests a tantalizing connection to schizophrenia, for which hallucinations and altered cognition are core symptoms. Schizophrenia also involves pathological neural pruning (Feinberg, 1982; Johnston, 2004; Sekar et al., 2016; Sellgren et al., 2019). Could pruning-induced degradation of internal predictive models explain some of the symptoms of this disease? The close relation between perception and reconstruction in DBMs and their hierarchical organization make them an interesting model candidate to further investigate hallucinations. Such modelling may provide theoretical insights into neurological disorders that involve aberrant pruning.

Apart from neurological disorders, understanding pruning is important for understanding learning and cognitive flexibility. In all of our experiments, the sensory encoding task was fixed. sentence broken: In reality, cognitive demands vary over the lifetime of an animal. Overzealous optimization to a fixed distribution could impair flexibility, a phenomenon that might relate to developmental critical periods and neurological disorders (Johnston, 2004). In biology, the brain must balance optimality against the need to maintain cognitive reserves to support ongoing learning and changing environments.

### 3.5 Biological plausibility of different pruning criteria

Pruning studies in artificial neural networks often estimate parameter importance with respect to the curvature of a cost function on network performance or discriminative power in supervised learning problems (e.g. LeCun et al., 1990; Hassibi et al., 1993; Sánchez-Gutiérrez et al., 2019). This requires access to global state, and is not applicable to biological neural networks. In contrast, our study examined unsupervised learning in sensory encoding models that seek to model the latent causes of sensory input (Hinton et al., 1995; Hinton, 2002; Hinton et al., 2006). We showed that an energy-based perspective on such encoders leads naturally to an implicit measure of parameter importance, and that single synapses could compute this measure.

We find that activity-driven pruning leads to better structural optimization than simply removing small weights. It identifies redundant units and protects computationally important synapses. Although parameter sensitivity correlates with weight values (Deistler et al., 2018), we show that activity-dependent pruning must consider more than the magnitude of a weight. Sampling the correlations between presynaptic and postsynaptic activity is one way to compute importance. We also show that importance can be estimated by combining weight magnitude with the firing rates of the presynaptic and postsynaptic neuron. This highlights that deceptively simple biological quantities can encode information-theoretic measures of synaptic importance.

Translated to biology, this implies that neurons and synapses are initially over-produced to allow for structural refinement of connectivity that depends on activity. In the encoding models explored here, neurons compete to explain the latent causes underlying sensory input. Pruning of overall unimportant neurons occurs if either set of incoming (dendritic) or outgoing (axonal) synapses is eliminated. To survive, neurons must maintain co-varying activity with both presynaptic inputs and post-axonal targets. Loss of either input drive, or output relevance, will lead to pruning.

### 3.6 Summary and outlook

Overall, we have shown that local activity-dependent synaptic pruning can solve the global problem of optimizing a network architecture. In contrast to pruning rules based on synaptic weights, our information-based procedure readily identified redundant neurons and led to more efficient and compact networks. The pruning strategy we outline uses quantities locally available to each synapse, and is biologically plausible. The artificial neural networks explored here are abstract, and whether an analogous process exists in biological neural networks remains to be explored. If analogous processes operate in biology, then a similar pruning procedure could optimize metabolic cost by eliminating redundant neurons.

## Author contributions

conceptualization C.S., M.E.R., M.H.H.; methodology, validation, and formal analysis C.S., M.H.H.; software, C.S.; writing–original draft preparation, C.S; writing– review, and editing, C.S., M.E.R., M.H.H.; supervision, project administration, and funding acquisition M.H.H.;

## Abbreviations

RBM: Restricted Boltzmann Machine
DBM: Deep Boltzmann Machine
FIM: Fisher Information Matrix
FI: Fisher Information

## 4 Materials and Methods

### 4.1 Datasets

Two different datasets of visual image patches were used to train and evaluate our models. First, 90, 000 circular patches of different size were randomly selected from images of the CIFAR-10 dataset (Krizhevsky et al., 2009) to mimic the encoding of visual scenes through receptive fields of retinal ganglion cells. Images were converted to gray scale and binarized according to their median pixel intensity. The procedure is illustrated in Figure 1 A.

Second, the MNIST handwritten digits dataset (LeCun et al., 2010) was used. Each square 28×28 images was cropped to 20×20 pixels by removing two pixels on each side. This resulted in a visible layer size of 400 units. The labeled images belonged to one of ten digit categories (0 − 9) and were divided in a training set of 60, 000 and a held-out test set of 10, 000 images. The gray scale images were binarized according to the mean pixel intensity in the training and test set, respectively.

### 4.2 Model definition and training

RBMs are generative stochastic encoder models consisting of *n* binary units (neurons) that are organized in two layers: a visible layer v that directly corresponds to a given input vector and a hidden (or latent) layer **h** which contain an encoding of the input data. Each visible layer neuron *v*_*i*_ has undirected weighted connections (synapses) to each hidden neuron *h*_*j*_ and vice versa, but neurons within a layer are not connected to each other (see Figure 1 A).

The energy function given in Equation 1 assigns energy to each visible layer pattern. It can be rewritten as a joint probability, which describes the system’s Boltzmann distribution at temperature *T* = 1:

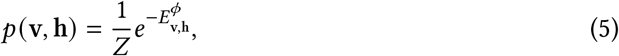

with *Z* being the partition function that sums over all possible configurations of visible and hidden neurons. By marginalizing and taking the logarithm, we get:

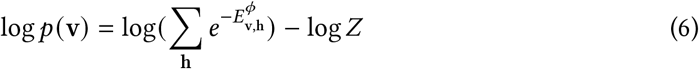

The training objective is to adjust the parameters *ϕ* such that the energy for a training pattern is lowered compared to the energies of competing patterns (Hinton, 2012). Lowering its energy translates to assigning a greater probability to that pattern. We maximize *p*(v) by maximizing the first term in Equation 6, i.e. the unnormalized log probability assigned to the training pattern **v**, and minimizing the second term. From the restricted connectivity it follows that the probability of a unit being in the on-state is conditionally independent from the states of units of the same layer, given a configuration of the other layer:

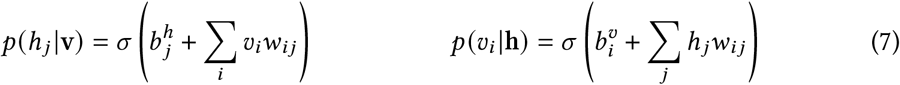

The positive-negative or wake-sleep algorithm (Hinton et al., 1995) exploits this conditional independence and is used to train RBMs. While the positive or wake phase corresponds to a forward-pass through the network (*p*(*h*_*j*_|**v**)), the model metaphorically dreams about *p*(*v*_*i*_|**h**) in the negative or sleep phase. Specifically, the visible layer is initialized to a random state and the layers are updated iteratively following alternating Gibbs sampling for *k* → ∞ times. The goal is for the final sample to originate from the equilibrium distribution of the model. *k*-step contrastive divergence (Hinton, 2002) accelerates the learning algorithm by effectively replacing the desired sample from the model distribution by a single reconstruction after *k* Gibbs steps. The most extreme case is one-step contrastive divergence, where the Gibbs chain is truncated after just one iteration of initializing the visible layer to a training example, sampling the hidden layer, and re-sampling the visible layer.

A DBM consists of a visible layer and multiple hidden layers. Analogous to Equation 1, the energy function of a two-layer Bernoulli-DBM is given by:

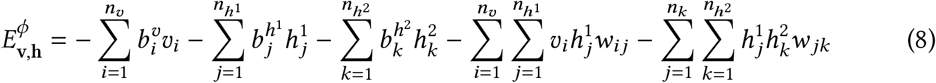

Our parameter set *ϕ* is thus augmented by an additional weight matrix 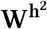 and another bias vector 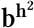. Learning in DBMs also follows the positive-negative algorithm, yet layer-wise: each layer of hidden units aims to find an appropriate representation of the distribution over the variables in its preceding layer in order to generate it. However, since contrastive divergence is too slow for training deeper networks (Hinton et al., 2006), a mean-field variational inference approach is used to train DBMs (Salakhutdinov and Larochelle, 2010).

### 4.3 Model fitting and sampling

All models were implemented in TensorFlow (Abadi et al., 2015) making use of open-source code for the implementation of RBMs and DBMs (https://github.com/yell/boltzmann-machines). Our pruning functionality was added in the forked repository (https://github.com/carolinscholl/PruningBMs). Gibbs sampling ran on NVIDIA GeForce GTX 980 or 1080 GPUs. For the layer states to be uncorrelated, a sample was stored after every 200th Gibbs step. The number of samples per layer was the same as the number of training instances.

#### 4.3.1 RBMs for CIFAR-10 patches

Single-layer RBMs were fit to CIFAR-10 patches using one-step contrastive divergence (Hinton, 2002). The radius of a circle determined the number of pixels and visible units (see Figure 1 A). Weights were initialized randomly from a Gaussian distribution with 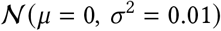. The hidden units outnumbered the visible units because we aimed for a sparse representation and uncorrelated hidden units. The initialization of hidden biases with −2 was another means to encourage sparseness in the firing rates of hidden neurons. Other than that, no sparsity target was defined. As recommended by (Hinton, 2012), visible biases were initialized with log [*p*(*v*_*i*_)/(1 − *p*(*v*_*i*_))], where *p*(*v*_*i*_) corresponds to the fraction of training instances where pixel *i* was in the on-state. No mini-batches were used, updates were applied after each training instance. RBMs were trained for 2 epochs, with a decaying learning rate from 0.1 to 0.01 and momentum of 0.9. Weights were not regularized to encourage a sloppy pattern in parameter sensitivities as is characteristic for biological networks (Daniels et al., 2008; Gutenkunst et al., 2007).

#### 4.3.2 DBMs for MNIST digits

The comparably large number of 400 pixels of each cropped MNIST image required their segmentation in receptive fields (see Figure 3 A). Hidden layer **h**^1^ had the same number of units as the visible layer **v**. Each unit from **h**^1^ was connected to a rectangle spanning a maximum number of 5 × 5 neighboring units from **v**. A stride of (1, 1) without zero-padding was used, so the receptive fields overlapped and were smaller at the borders of the image. The use of receptive field reduced the number of weights connecting **v** and **h**^1^ from originally 400 × 400 = 160, 000 to 8, 836. The fully connected hidden layer **h**^2^ then combined the receptive field encodings of parts of the image into a latent representation of the full image.

The DBM was built from two individual RBMs that were pre-trained for 20 epochs with one-step contrastive divergence. After fitting the first RBM with receptive fields, its hidden units were sampled with v clamped to an input vector at a time. The resulting binary hidden layer vectors served as training data for the second RBM with 676 fully connected hidden units. Weights for both RBMs were initialized randomly from a Gaussian distribution with 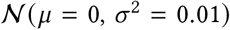. Visible biases were initialized with −1, hidden biases with −2. Neither sparsity targets nor costs were defined. Momentum was set to 0.5 for the first 5 epochs and then increased to 0.9 as recommended by (Hinton, 2012). The learning rates decreased on a logarithmic scale between epochs, starting from 0.01 for the first RBM, and from 0.1 for the second RBM, to 0.0001 in their final epochs of training. No mini-batches or weight regularization methods were used.

When stacking the two RBMs, the hidden layer of the first RBM and the visible layer of the second RBM were merged by averaging their biases. The resulting DBM with two hidden layers was trained jointly for 20 epochs following a mean-field variational inference approach using persistent contrastive divergence with 100 persistent Markov chains (Salakhutdinov and Larochelle, 2010). A maximum value of 6 was defined for the norm of the weight matrix to prevent extreme values. The hidden unit firing probabilities decayed at a rate of 0.8. Neither a sparsity target nor weight regularization were applied and parameters were updated immediately after presenting a training image.

### 4.4 Pruning criteria

We compared six different weight pruning criteria throughout our simulations, three of which targeted the removal of weights that carried low FI. For small RBMs, computing the full FIM and its eigenvectors was feasible, using code from github.com/martinosorb/rbm_utils. The parameter-specific entries of the first eigenvector were then used as indicators of parameter importance. For DBMs we only used two of the three FI pruning criteria: Variance FI refers to estimating the weight-specific FIM diagonal entries by computing (4) based on samples from all layers of the models. Heuristic FI does not directly track the firing rates, but uses the mean rates instead (see Appendix A.2). For the DBM, the FIM diagonal was computed layer-wise assuming weakly correlated hidden units. To examine the relevance of high FI parameters, we also deleted the *most* important weights according to the variance estimate of FI, which we refer to as Anti-FI pruning. For heuristic FI, we replaced the second term in Equation 4 with a mean-field estimate of the covariance of units firing (Eq. A2). When we used the weight magnitude as a proxy of importance, we deleted weights with lowest absolute value. As a baseline, we also removed a randomly chosen sample of weights.

#### 4.4.1 Pruning procedure

To simulate synaptic pruning, weight importance was estimated according to the selected pruning criterion. According to this criterion, a threshold was set at a pre-specified percentile. All weights whose estimated importance did not meet this threshold were removed from the network by fixing their values to zero. For RBMs trained on CIFAR-10 patches, the threshold corresponded to the 50^th^ percentile of weight importance. For DBMs trained on MNIST, the threshold corresponded to the 10^th^ percentile of weight importance. If all incoming weights to a hidden unit were pruned, the unit was removed from the network. In DBMs, a hidden unit was also deleted if all its outgoing weights were pruned. After pruning, the model was re-initialized with unaltered values of the remaining parameters for the next pruning iteration. The RBMs fit to CIFAR-10 patches were retrained for 2 epochs. The retraining period for DBMs was shortened from 20 to 10 epochs after pruning. This pruning procedure was repeated three times for experiments with RBMs and ten times for experiments with DBMs.

For the experiments on MNIST, we also examined a full retraining phase after the final iteration of pruning. This included a random re-initialization of parameters and individual pre-training of the two RBMs before stacking and retraining the full DBM. All training hyper-parameters were the same as for the original DBM (see Section 4.3.2).

### 4.5 Evaluation

Each experiment started by fitting a sufficiently large model to the data. While the visible layer size was determined by the number of pixels in each training image, the (final) hidden layer was set to be larger. The resulting over-parameterized model was iteratively pruned, while its fit was monitored.

For small RBMs that were trained on CIFAR-10 patches, we evaluated a model’s generative performance by comparing its generated visible layer samples to the frequencies of patterns occurring in the training data. For DBMs that were trained on MNIST digits, we made use of the labeled data to evaluate both the encoding and generative performance.

First, we considered the encodings of the data in the final hidden layer of the DBM. While the visible layer was clamped to one training instance at a time, the hidden unit activations were sampled. We expect these latent representations to comprise a more useful training set for the classifier than the raw images. The resulting set of 60,000 final hidden layer encodings was used to train a multinomial logistic regression classifier, which had to distinguish between the 10 digit categories. We refer to the classification accuracy of this classifier built on top of the final hidden layer as the encoding quality of the model.

Second, we evaluated the patterns generated by the DBM. Since Boltzmann machines try to approximate the data distribution with their model distribution, these generated patterns should ideally resemble digits. Thus, we trained a multinomial logistic regression classifier on the 60,000 raw MNIST images. After training, this classifier received patterns generated by the DBM in order to classify them. For each of the ten digit classes, it returned a probability of the current sample belonging into it. The argmax over this probability distribution was used to assign the class. The average of the winning class probabilities was used as a confidence score. It served as a measure of digit quality. Furthermore, the distribution of assigned classes served as a measure of digit diversity. Specifically, we counted the samples that were assigned to each of the 10 digit classes and computed their fraction of generated samples. If the generated samples were completely balanced across classes, we would expect each digit to be generated in 10% of cases.

Moreover, the encoding performance was compared to the accuracy of a classifier trained on the raw digits. The quality of generated digits was compared to the quality of the 10,000 held-out MNIST test digits and to the quality of random patterns, using the same classifier.

For the DBMs we also evaluated the generative performance of a fully retrained pruned model after the final pruning iteration. For this, we generated 60,000 visible samples from these pruned retrained models as before. As explained above, we let the classifier trained on the original MNIST dataset classify these samples. The entropy over the resulting class counts served as another metric of digit diversity. To get a confidence bound on the estimated entropy, we performed bootstrapping. We took 10,000 samples of fixed size *k*, with *k* ranging from 100 to 80,000. We then computed the mean entropy and its standard deviation per sample size.

## A Local estimates of Fisher Information

## A.1 Variance estimate of Fisher Information

Parameter importance is reflected in the curvature of the energy landscape of an RBM when slightly changing two parameters. Computing this for each parameter pair leads to the FIM (see Equation 2), where *F*_*i*_ stands for the Fisher information of the considered couple (*ϕ*_*i*_, *ϕ*_*j*_). The entries of the FIM thus have the form (Rule et al., 2020):

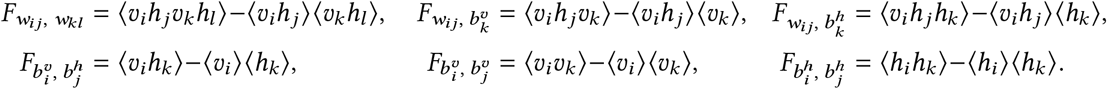

The diagonal of the FIM corresponds to evaluating changes in the energy landscape of the model when perturbing just one parameter. The importance of a weight thus simplifies to the average coincident firing of pre- and postsynaptic neurons. The importance of a bias value is estimated by the variance of a neuron’s firing.

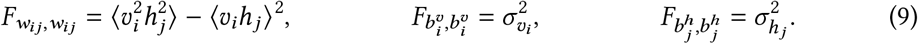

We consider both of these estimates to be locally available statistics to the individual neuron.

## A.2 Heuristic estimate of Fisher Information

In this section, we derive a mean-field approximation estimate of 〈*v*_*i*_*h*_*j*_〉 in terms of a small deviation from the case where presynpatic and postsynaptic activity are statistically independent, 〈*v*_*i*_*h*_*j*_〉 ≈ 〈*v*_*i*_〉 〈*h*_*j*_〉 that accounts for correlated activity introduced by the synapse between neurons *i* and *j*. Starting from Equation 4, we get

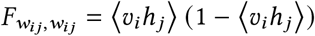

This can be computed if there is a local mechanism for tracking and storing the correlation of pre- and postsynaptic firing rates 〈*v*_*i*_*h*_*j*_〉. This expectation is closely related to more readily available local statistics, like the mean rates and weight magnitude. Since *v*_*i*_ and *h*_*j*_ binary in {0, 1}, the expectation 〈*v*_*i*_*h*_*j*_〉 amounts to estimating the probability that *v*_*i*_ and *h*_*j*_ are simultaneously 1, i.e. *p*(*v*_*i*_=1, *h*_*j*_ =1). One can express this in terms of a mean rate and the conditional activation of presynaptic neuron given a postsynaptic spike, using the chain rule of conditional probability:

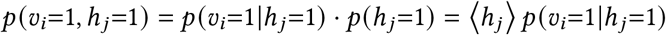

If the hidden layer size is large and the hidden units are independent, we may approximate the activity of all *other* hidden units apart from a particular *h*_*j*_ using mean-field. The activation of a visible unit is given by Equation 7. We replace the hidden units by their mean firing rate:

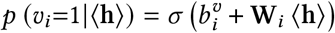

To estimate the contribution of all units *except h*_*j*_, one computes:

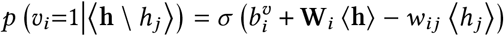

To get the mean-field activation assuming *h*_*j*_ = 1,

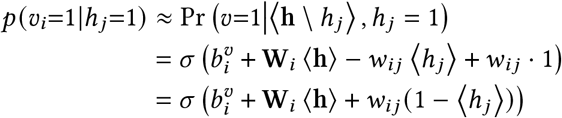

We can obtain an alternative (and, empirically: more accurate) mean-field approximation using the average firing rate of the visible unit, 〈*v*_*i*_〉. Given this mean rate, we can estimate the activation as *σ*^−1^(〈*v*_*i*_〉). This estimate can replace the 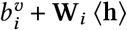 terms, leading to:

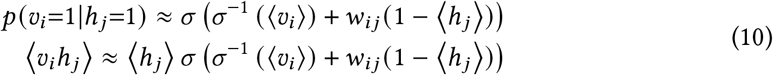

This approximation is ad-hoc, but captures an important intuition: the magnitude of synaptic weights themselves provides a useful proxy for computing pre-post correlations, and therefore estimating a synapse’s importance in the network.

